# Unique axon-to-soma signaling pathways mediate dendritic spine loss and hyper-excitability post-axotomy

**DOI:** 10.1101/642207

**Authors:** Tharkika Nagendran, Anne Marion Taylor

## Abstract

Axon damage may cause axon regeneration, retrograde synapse loss, and hyper-excitability, all of which affect recovery following acquired brain injury. While axon regeneration is studied extensively, less is known about signaling mediating retrograde synapse loss and hyper-excitability, especially in long projection pyramidal neurons. To investigate intrinsic injury signaling within neurons, we use an *in vitro* microfluidic platform that models dendritic spine loss and delayed hyper-excitability following remote axon injury. Our data show that sodium influx and reversal of sodium calcium exchangers (NCXs) at the site of axotomy, mediate dendritic spine loss following axotomy. In contrast, sodium influx and NCX reversal alone are insufficient to cause retrograde hyper-excitability. We found that calcium release from axonal ER is critical for the induction of hyper-excitability and inhibition loss. These data suggest that synapse loss and hyper-excitability are uncoupled responses following axon injury. Further, axonal ER may play a critical and underappreciated role in mediating retrograde hyper-excitability within the CNS.

## Introduction

Acute neural injuries (e.g., stroke, traumatic brain injury, and spinal cord injury) cause profound axon damage. Axon damage triggers an intra-cellular signaling cascade to effect neuronal injury responses, including axon regeneration, retrograde synapse loss, and hyper-excitability. These downstream responses are critical for recovery following injury. Yet, the intrinsic neuronal signaling mechanisms mediating retrograde synapse loss and hyper-excitability, in particular, remain poorly understood.

Axon injury induces differential gene expression and transcription within the soma, requiring long range signaling from the site of injury to the nucleus (Rishal and Fainzilber 2014; Nagendran et al. 2017). Breach of the axonal membrane following axon injury causes an influx of ions, including calcium and sodium, into the intra-axonal space. The increase in sodium ions through voltage-gated sodium channels causes reversal of sodium-calcium exchangers (NCXs) located on the plasma membrane, mitochondria and ER, thus enhancing local intra-axonal calcium levels (Persson et al. 2013). Calcium release from smooth ER within the axon may also potentiate axon-to-soma signaling (Cho et al. 2013; Sun et al. 2014). Axon damage of mouse peripheral sensory neurons was reduced with blockade of both sodium channels and the reverse mode of NCX (Persson et al. 2013), supporting the critical role of NCXs in retrograde injury signaling. Whether local sodium influx and reversal of NCX during axon damage are needed to transmit signals to the soma to cause dendritic spine loss and hyper-excitability remains unknown.

Hippocampal cultures grown within a compartmentalized microfluidic platform provide an injury model system to investigate intrinsic neuronal injury response. These devices guide axonal growth of pyramidal cells, through a microgroove-embedded barrier region of almost 1 mm into an isolated axonal compartment. Because of this barrier region, axons can be injured precisely without mechanically disrupting somatodendritic regions and soluble microenvironments established for experimental purposes. Axotomy performed within compartmentalized platforms produced several characteristic injury responses well described *in vivo*, including rapid expression of the immediate early gene *c-fos* (Taylor et al. 2005) and reduced expression of netrin-1 one day following axon damage. Evidence of ER changes within the soma, called chromatolysis, occurs in axotomized neurons in vitro (McIlwain and Hoke 2005; Nagendran et al. 2017). Axotomized neurons within microfluidic chambers also showed that dendritic spine loss (Nagendran et al., 2017)(Gao et al. 2011; Ghosh et al. 2012) and hyper-excitability (Frost et al. 2015; Nagendran et al. 2017) occur, providing an experimentally tractable model to study initiation and progression of these neuronal injury responses. Axotomy within these platforms produced a delayed enhancement in neuronal hyper-excitability, one day following dendritic spine loss. The extent to which both dendritic spine loss and hyper-excitability use the same retrograde signaling mechanisms is unclear.

## Results

### Sodium influx and reversal of sodium calcium exchangers induce retrograde spine loss

To selectively injure axons of hippocampal neurons > 900 μm from somata, we used compartmentalized microfluidic devices (**Fig. 1a**). Primarily axons of pyramidal neurons are guided via microgrooves into a separated axon compartment where they are injured and various pharmacological treatments can be restricted to isolated axons. We previously found that blocking local activity at the site of injury using this method prevented dendritic spine loss 24 h post-axotomy (Nagendran et al. 2017), suggesting calcium and sodium influx at the site of injury are key mediators of this neuronal injury response. Reversal of NCX at the site of injury may play a key role, causing this massive local influx of calcium. To determine whether local NCX activation is required to trigger retrograde synapse loss, we performed axotomy within microfluidic devices but in the presence of the reversible NCX blocker applied specifically to the axonal compartment where axotomy was performed (**Fig. 1b**). We quantified spine density using repeated live imaging and found that in the presence of the NCX blocker, KB-R7943, spine loss was completely prevented (**Fig. 1b,c**). Controls treated with vehicle had significantly fewer spines following axotomy (**Fig. 1b,c**).

**Figure 1:**
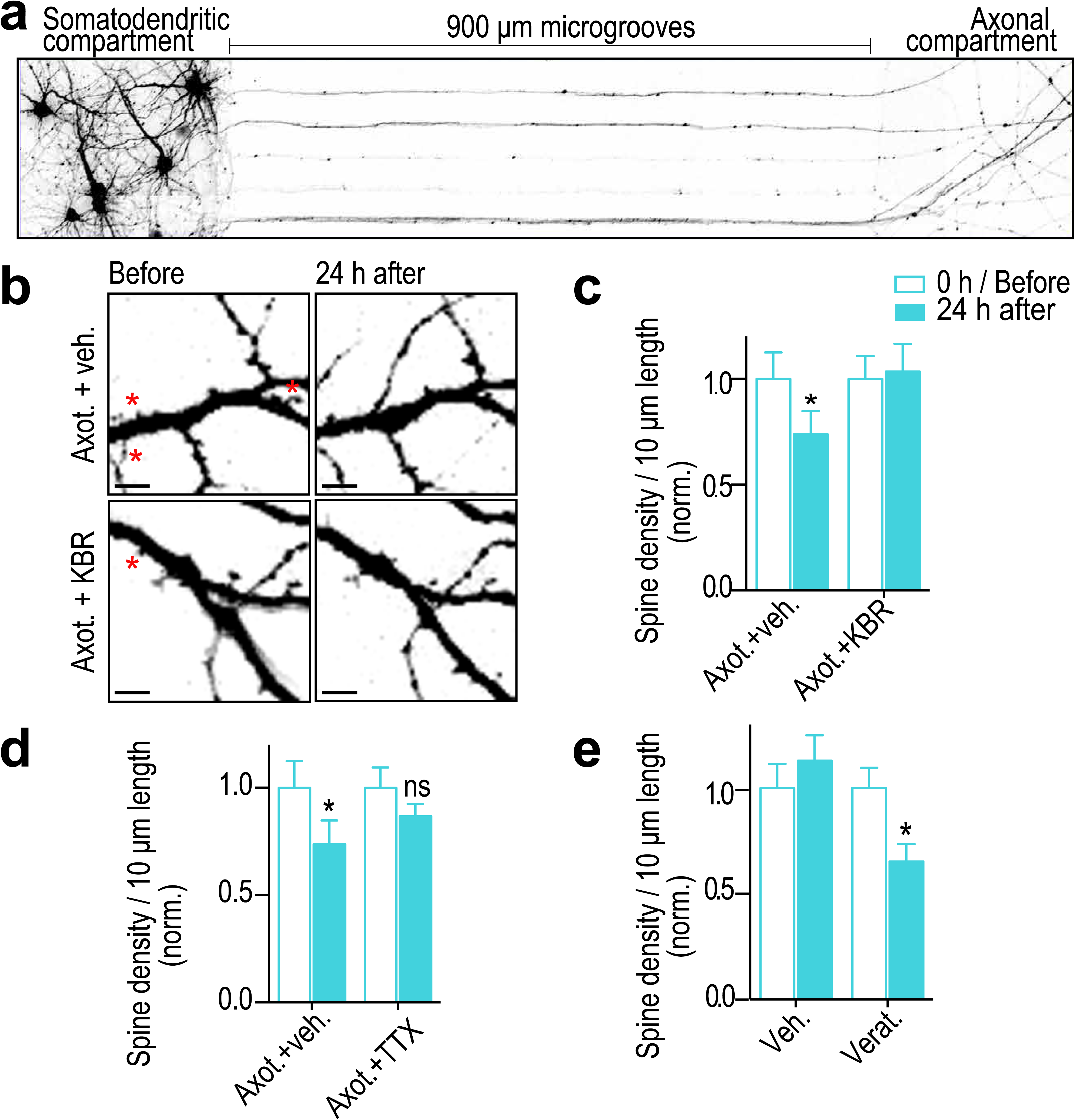
Blocking sodium channels and reverse mode of NCX at the site of injury prevents axotomy-induced spine loss. **(a)** Inverted gray scale images of 15 DIV rat hippocampal neurons cultured within microfluidic devices that were retrogradely labeled using G-deleted mCherry rabies virus added to axonal compartment. **(b)** Representative images of dendrites before and 24 h after axotomy treated with either vehicle or reversible NCX blocker (KB-R7943; 10 μM). Astericks indicate eliminated spines. Scale bars, 5 μm. **(c, d)** Quantification of dendritic spine density before and 24 h after axotomy with application of either vehicle or KBR or TTX (1 μM) that was applied only to the axonal compartment for 1 h during injury. **(e)** Quantification of spine density before and 24 h after treatment of axonal compartment with either vehicle or sodium channel activator (veratridine, 10 μM) for 10 min in the absence of injury. n=15 dendrites for each condition over 2 independent experiments. Paired two-tailed t-test, *p ≤ 0.05. *Error bars*, s.e.m.

Sodium influx at the site of injury may trigger reversal of NCX needed for dendritic spine loss. Thus, we blocked sodium channels using tetrodotoxin (TTX) at the site of injury and quantified dendritic spine density changes. As expected, blocking sodium channels prevented a significant reduction in dendritic spine density (**Fig. 1d**). Further, application of a potent activator of voltage gated sodium channels, veratridine, at distal axons and in the absence of injury led to a significant decrease in retrograde spine density 24 h post treatment. Together, these results show that sodium influx and reversal of NCXs at the site of injury trigger retrograde spine loss.

### Calcium influx via reversal of NCX is not required to induce retrograde hyperexcitability post-axotomy

We next tested whether retrograde hyper-excitability is triggered via reversal of NCXs. To examine retrograde hyperexcitability, we used FM dyes to quantify synaptic vesicle release dynamics as performed previously (Nagendran et al. 2017). This is an unbiased approach to measure synaptic vesicle release in response to field stimulation. Previously published data showed that synaptic vesicle release increases significantly 48 h post axotomy, analyzed via FM release curves and frequency of miniature excitatory postsynaptic currents (Nagendran et al. 2017). As a control we repeated these experiments and also tested whether blocking NCX reversal at the site of injury prevents the increase in hyper-excitability (**Fig. 2a,b**). Surprisingly, we found that blocking NCX reversal at the site of injury did not prevent the increase in release rate due to axotomy. In fact, the KBR-treated cultures were indistinguishable from the vehicle controls. We performed a similar experiment using TTX (**Fig. 2c**). Again, this did not prevent axotomy-induced hyper-excitability. In fact, the increase in release rate was exacerbated with the TTX treatment. We next tested whether veratridine in the absence of injury would cause hyper-excitability (**Fig. 2d**). Consistent with our preceding data, the FM release curves were indistinguishable from vehicle controls. These results suggest that retrograde hyper-excitability involves a unique triggering mechanism independent from retrograde spine loss signaling following axotomy.

**Figure 2:**
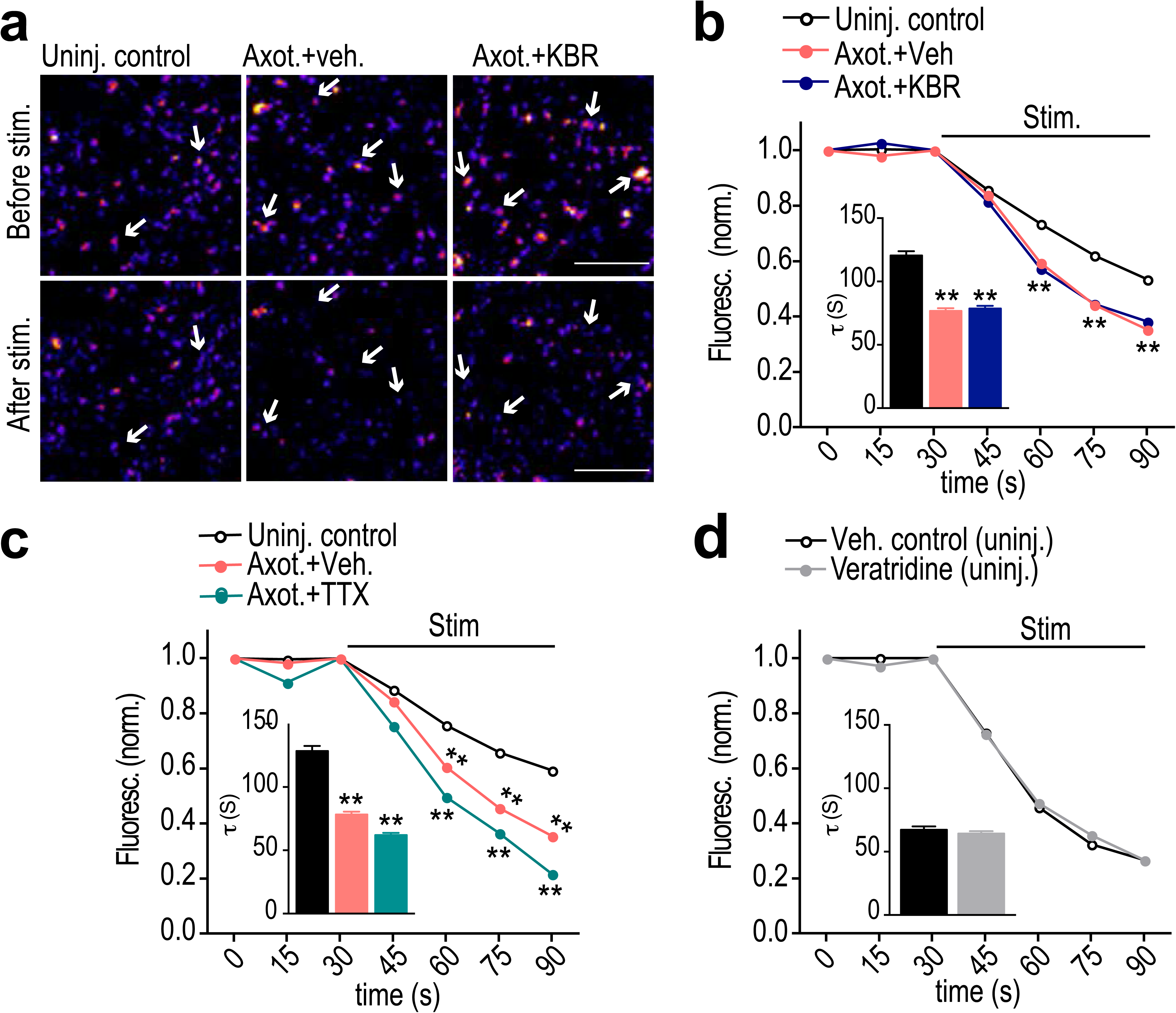
Calcium influx via reversal of NCX is not required to induce retrograde hyperexcitability post-axotomy. Representative images of presynaptic terminals labeled with FM5-95 (FM puncta) before and after field stimulation within the somatodendritic compartment of microfluidic chambers. Color look up table ‘Fire’. Scale bars, 10 μm. **(b, c)** FM unloading curves at 48 h following application of KBR or TTX **(c)** to axonal compartment for 1 h during axotomy. Uninjured control: n= 782 puncta; axoyomy: n= 1037 puncta; Axot. KBR: n=1012 puncta; Axot. TTX: 1415 puncta. **(d)** FM unloading curves at 48 h following 10 min application of veratridine to axonal compartment without injury. Control (DMSO): n=711 puncta; veratridine: n=1273 puncta. Two-way ANOVA, Bonferroni post hoc test. Inset in *b, c, d* shows FM decay time constant (*τ*) for puncta with *τ* <360s (control, *n*=654; axotomy, *n*= 977; Axot. KBR: n=945 puncta; Axot. TTX: 1321 puncta). *(d)* control, *n*=634; veratridine, *n*=1177. Unpaired two-tailed *t*-test. 5-6 chambers for each condition over 3 independent experiments. \*\**p*<0.001. *Error bars*, s.e.m.

### Retrograde hyperexcitability post-axotomy requires calcium release from ER intracellular stores

Calcium-mediated signaling is expected to be a critical factor in triggering retrograde hyper-excitability following injury. To further investigate the role of calcium, we applied low calcium and TTX solution within the axonal compartment at the time of injury. Surprisingly, this treatment did not alter the axotomy induced FM release kinetics compared with injured vehicle control (**Fig. 3a,b**). Intracellular calcium stores may also play a critical role in axon-to-soma injury signaling, thus we next applied the low calcium/TTX solution together with blockers of ER calcium release from ryanodine (Dantrolene) and IP3 receptors (Xestospongin C). Blocking calcium release from ER normalized the synaptic vesicle release rate to uninjured control levels (**Fig. 3c,d**).

**Figure 3:**
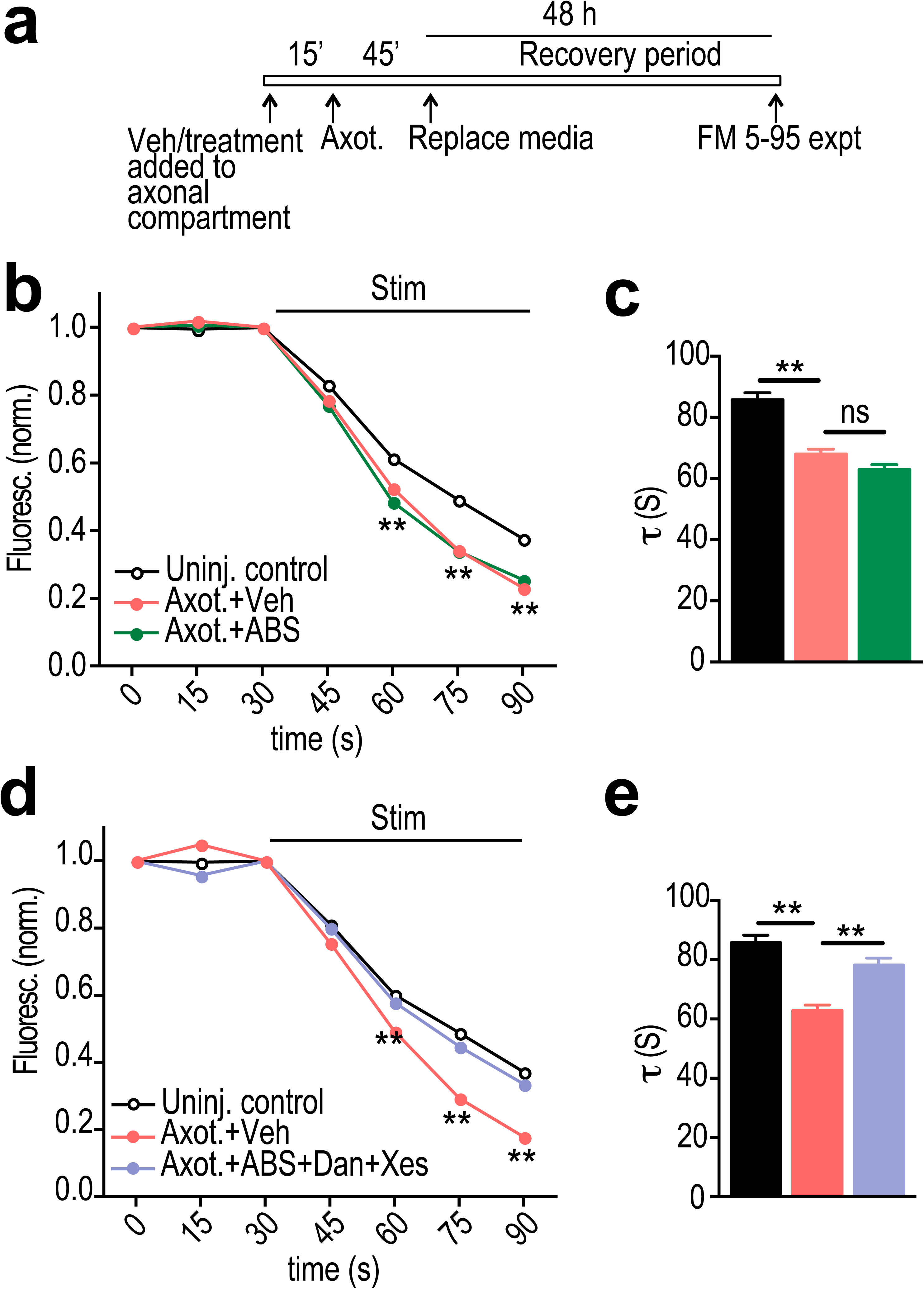
ER calcium channel activation at the site of injury is required for retrograde presynaptic hyper-excitability. **(a)** Experimental timeline for treatment and imaging within microfluidic chambers. **(b)** FM unloading curves at 48 h following application of vehicle or local activity blockade solution (ABS) to axons for 1 h during axotomy. Uninjured control: n= 1135 puncta; Axot. n=1834 puncta; Axot. ABS: n=1590 puncta; 8 chambers per condition over 4 experiments. Two-way ANOVA, Bonferroni post hoc test. **(c)** FM decay time constant (*τ*) for puncta with *τ* = <360s (48h control, *n*=1044; 48h axotomy, *n*=1686; 48h Axot. ABS, *n*=1478). Unpaired two-tailed *t*-test. Each condition includes 8 chambers over 4 experiments. **(d)** FM unloading following 48 h after application of vehicle or local activity blockade solution (ABS) supplemented with ryanodine receptor (Dantrolene, 20μm) and IP3 receptor (Xestospongin C, 1μm) blockers to axonal compartment with 15 min pre-treatment and 45 min treatment post-axotomy. Uninjured control: n= 974 puncta; Axot. n=1239 puncta; Axot. ABS + Dan + Xes: 1124 puncta; 5 to 6 chambers per condition over 3 experiments. Two-way ANOVA, Bonferroni post hoc test. **(e)** FM decay time constant (*τ*) for puncta with *τ* = <360s (48h control, *n*=892; 48h axotomy, *n*=1128; 48h Axot. ABS + Dan + Xes, *n*=1025). Unpaired two-tailed *t*-test. Each condition includes 5–6 chambers over 3 experiments. \*\**p*<0.001. *Error bars*, s.e.m.

We previously found that the increase in FM release rate following injury coincided with loss of inhibitory terminals onto injured neurons. Loss of inhibition is a cause of hyper-excitability. As confirmation that low calcium/TTX did not substantially influence axotomy-induced disinhibition, we quantified the number of vGAT immunolabeled puncta colocalized with the dendritic arbor of labeled axotomized neurons in the presence of low Calcium/TTX solution (**Fig. 4a**). Application of this solution did not prevent the loss of inhibitory terminals. In contrast application of this solution with Dant and Xesto to block ER-dependent calcium release caused injured neurons to retain inhibitory terminals (**Fig. 4b**). These data show that ER provides a critical source of calcium needed to propagate the transsynaptic change of altering synaptic vesicle release and inhibitory terminal loss.

**Figure 4:**
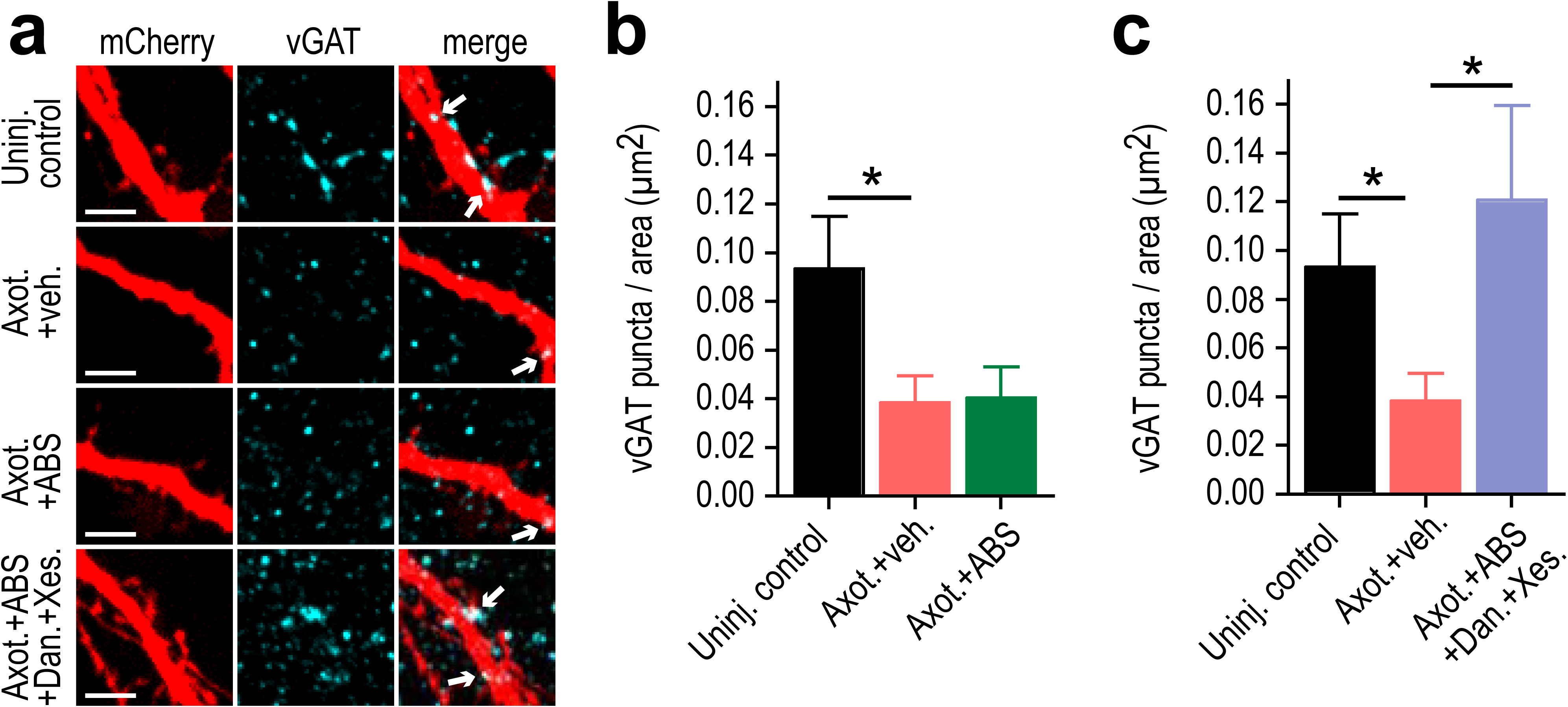
ER calcium channel activation at the site of injury is required for retrograde inhibitory terminal loss post-axotomy. **(a)** Representative mCherry labeled dendritic segments immunostained for inhibitory presynaptic marker, vGAT, at 48 h following application of vehicle or local activity blockade solution (ABS) with or without Dantrolene and Xestospongin C to axons for 1 h during axotomy. Arrows in the merge image indicate the localization of vGAT puncta (cyan) on mcherry labeled dendrite (red). Scale bar, 5 μm. **(b,c)** Number of vGAT puncta per neuron area (uninjured control, axotomy or axotomy + ABS or axotomy + ABS + Dan + Xes) at 15-16 DIV. *n*=6-8 neurons; 3 chambers per condition over 2 independent experiments. Error bars, SEM. *P<0.05.

### ER-dependent changes following axotomy

Our data suggests that ER plays a critical role in trans-synaptic injury signaling following axotomy. To investigate further how somatodendritic ER changes following axotomy, we measured the changes in the ER stress marker, Bip, using immunolabeling (**Fig. 5a,b**). Significant Bip accumulation occurred within the soma at 12 and 24 h post-axotomy. We also found significant upregulation of the ER calcium pump, SERCA2, within the somatodendritic compartment following axotomy (**Fig. 5a,c**).

**Figure 5.**
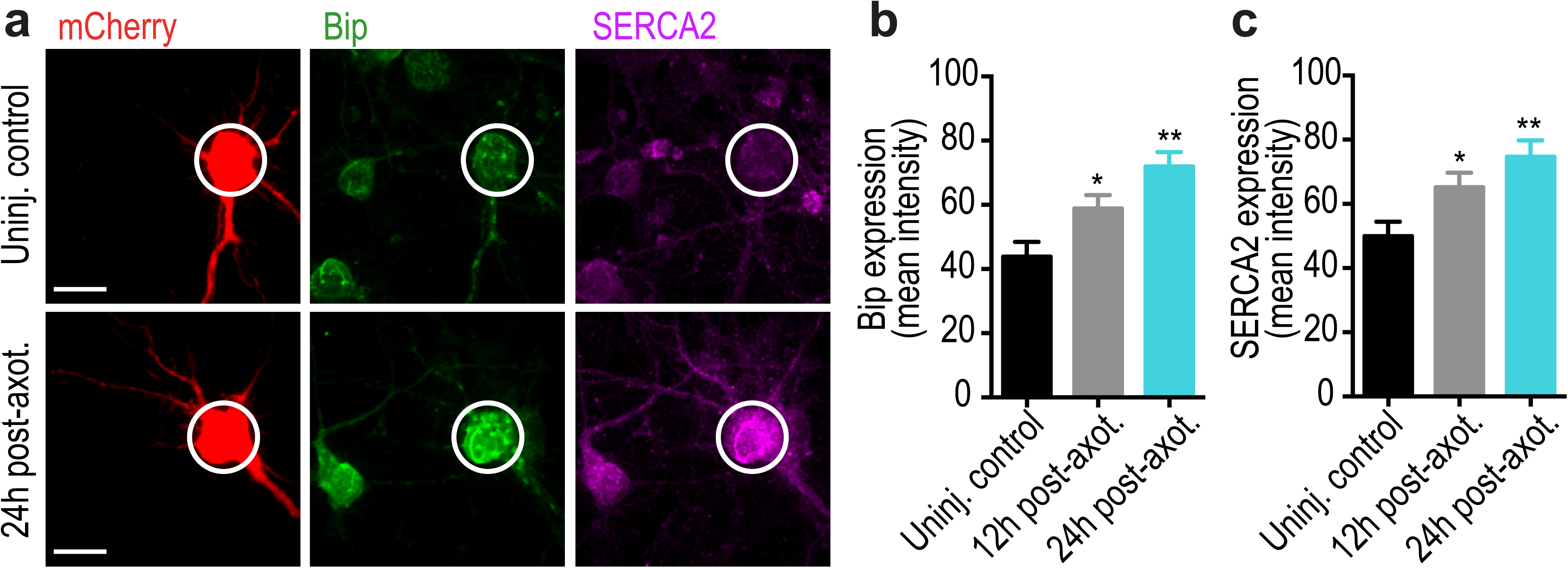
Axotomy induces somatic ER stress within our microfluidic model. **(a)** Representative images of retrograde labeled mCherry neurons co-immunostained with ER stress markers, GRP78/Bip and SERCA2. Scale bar, 10 μm. White circles highlight the soma of axotomized and uninjured control neurons. Quantification of somatic Bip **(b)** and SERCA2 **(c)** expression at 12 and 24 h post-axotomy, indicative of somatic ER stress. N=18 neurons; 2 chambers per condition. Error bars, SEM. *P<0.05, **p<0.005.

## Methods and Materials

### Microfluidic chambers

Poly (dimethylsiloxane) (PDMS) was molded onto a SU-8 master with 900 μm long, 3–4 μm tall and 7.5–8 μm wide microgrooves as previously described (Taylor et al. 2003; Taylor et al. 2005). Chambers were sterilized in 70% ethanol and placed onto 500–550 kDa Poly-D-Lysine (BD Biosciences) coated sterile German glass coverslips.

### Hippocampal cultures

Animal procedures were approved by the University of North Carolina at Chapel Hill Institutional Animal Care and Use Committee (IACUC). Sprague Dawley rat embryos (E18-E19) were used to prepare dissociated hippocampal cultures as previously described (Nagendran et al. 2017). Hippocampal cells were dissociated in neuron culture media i.e., neurobasal media (Invitrogen) supplemented with 1 X B27 (Invitrogen), 1 × Antibiotic-antimycotic (Invitrogen), and 1 X Glutamax (Invitrogen). Approximately ∼90,000 cells were plated into the somatodendritic compartment of the chamber. After 5–7 days of culture in microfluidic chambers, axons extended into the adjacent axonal compartment.

### Retrograde labeling and axotomy

Neurons were retrogradely labeled with G-deleted Rabies-mCherry or eGFP virus between 11 and 13 days *in vitro* (Wickersham et al. 2007) (Salk Institute; 1 × 10^5^ viral units) as previously described (Nagendran et al. 2017). G-deleted Rabies-mCherry or eGFP virus diluted in 200 μl neuron culture media was added to the axonal compartment of each chamber. After 2 h incubation with virus at 37 °C remove virus containing media from the axonal compartment. Saved conditioned media was added back to the axonal compartments following two washes with fresh culture media. Chambers were maintained in 37 °C incubator for ∼48 h until fluorescence expression was visible. Axotomy was performed between 13 and 15 days in vitro (DIV) as previously described (Nagendran et al. 2017).

### Immunocytochemistry

Neuronal cultures were fixed with 4% PFA and permeabilized in 0.25% Triton X-100. Coverslips were blocked in 10% normal goat serum for 15 min each and incubated with anti-vGLUT1 (1:100; NeuroMab, clone N28/9, #75-066), anti-vGAT (1:1000; Synaptic Systems #131 003), anti-SERCA (1:500; Abcam # ab2817), and anti-GRP78 Bip (1:200; Abcam # ab21685) primary antibodies in 1% blocking solution for overnight at 4 °C. Coverslips were then incubated with goat anti-rabbit or goat anti-mouse secondary antibodies conjugated to Alexa-fluorophores (1:1000; Invitrogen) for 1 h at RT.

### FM dye experiments and analysis

48 h (15 DIV) post-axotomy cultures were loaded with lipophilic dye N-(3-trimethylammoniumpropyl) -4-(6-(4-(diethylamino) phenyl) hexatrienyl)pyridinium dibromide (FM 5–95; Invitrogen) using KCl mediated depolarization as described previously (Taylor et al. 2013). Microfluidic chambers were stimulated using extracellular electrodes by placing a positive and negative electrode in each well of the somatodendritic compartment. Electrical stimulation was provided by an AD Instruments 2 Channel Stimulus Generator (STG4002) in current mode with an asymmetric waveform (−480 μA for 1ms and + 1600 μA for 0.3 ms) for ∼ 1 min at 20 hz for 600 pulses. The FM 5–95 imaging was performed using a spinning disk confocal imaging system as previously described in Taylor et al. 2013. Z-stacks (31 slices) were captured every 15 s during the baseline (1 min), stimulation (1 min), and after stimulation (2 min) periods. At least 3 baseline images were acquired before electrical stimulation. Sum projected confocal z-stack were converted to 8-bit images and registered using TurboReg, an Image J plugin. We background subtracted the image stack using the image 3 min after stimulation began as described in Nagendran et al. 2017. Briefly, image stacks were thresholded to a pixel value of 15. FM puncta between 0.4 to 10 μm^2^ were analyzed. We measured the intensity of each punctum in the whole field throughout all time series.We normalized fluorescence intensity of each puncta to the frame before stimulation. Puncta with >5% unloading after 1 min were used in the analysis as unloaded puncta. Time constants were estimated by curve fitting unloading kinetics to a single exponential decay function (Taylor et al. 2013). Curve fitting was done in MATLAB and FM puncta with time constants longer than 3 min were excluded from the analysis.

### Drug treatments

KB-R7943 (Tocris Bioscience # 1244) was suspended in DMSO and applied to the axonal compartment at a final concentration of 10 μM for 1 h during axotomy (including 15 min pre-incubation before axotomy). Tetrodotoxin citrate (TTX; Tocris Bioscience #1078) was suspended in HBS and applied to the axonal compartment at a final concentration of 1 μM for 1 h during axotomy (beginning 15 min prior to axotomy). Veratridine (Tocris Bioscience #2918) was suspended in DMSO and applied to the axonal compartment at a final concentration of 10 μM for 10 min in the absence of axotomy/injury. Local ABS, which includes low-Ca^2+^, high-Mg^2+^, and TTX (0.5mM CaCl_2_, 10mM MgCl_2_, 1 μM TTX) was applied solely to the axonal compartment for 1 h during axotomy (15 min prior and 45 min after axotomy). Dantrolene (Tocris Bioscience #0507) and (−)-Xestospongin C (Tocris Bioscience #1280), stock concentrations prepared in DMSO, were diluted in ABS solution and added to axonal compartment at a final concentration of 20 μM and 1 μM respectively for 1 h during axotomy (with 15 min pre-treatment and 45 min treatment post-axotomy). DMSO or HBS was used as vehicles. Media stored from the axonal compartment prior to treatment was added back to the axonal compartment after treatment and washes with pre-warmed fresh neuron culture media.

### Microscopy and image analyses

FM and fixed imaging was performed using CSU-X1 (Yokogawa) spinning disk confocal imaging unit configured for an Olympus IX81 microscope (Andor Revolution XD). Excitation for the spinning disk confocal imaging system was provided by 405 nm, 488 nm, 561 nm, and/or 640 nm lasers. The following bandpass emission filters (BrightLine, Semrock) were used for the spinning disk: 447/60 nm (TRF447-060), 525/30 nm (TRF525-030), 607/36 nm (TR-F607-036), and 685/40 nm (TR-F685-040). Zeiss LSM 780 (63 × 1.4 NA or 40 × 1.4 NA oil immersion objective) or the spinning disk system above (60 × 1.3 NA silicon oil immersion objective) was used to capture high-resolution images of mCherry or eGFP labeled live neurons as previously described (Nagendran et al. 2017).

For FM imaging, the spinning disk confocal imaging system was used with excitation at 561 nm and the 685/40 nm emission filter. We used 2 × 2 binning to reduce the laser intensity and acquisition time for each frame; each z-stack was obtained in ∼5 s. For dendritic spine analysis, before and 24 h post-axotomy confocal z-stack images were captured from live cultures to create montages of neurons extending axons into the axonal compartment. Calibrated z-stack montages were analyzed for all dendrite and spine parameters. Primary dendrites were traced using the semiautomatic neurite tracing tool, Neuron J (Meijering et al. 2004). The number of spines on all primary dendrites of each neuron was manually counted and spine density was calculated for 10 μm length of dendrite as [(# of spines/dendrite length) x 10] (Nagendran et al. 2017).

### Statistical Analysis

Statistics was performed using GraphPad Prism 6. For sample size, p-value, and statistical test (see figure legends).

## 1 Discussion

Hyper-excitability following acquired brain injury leads to long-term effects, such as persistent seizures, chronic pain, and spasticity. Intrinsic injury signalling within damaged neurons likely plays a key role in induced hyper-excitability. Using our *in vitro* microfluidic model, our data suggest that ER-dependent calcium release within the axon at the site of injury is a critical signalling component that triggers changes in retrograde presynaptic neurotransmitter release rate. Blocking axotomy-induced dendritic spine loss through local activity blockade does not prevent hyper-excitability, suggesting that hyper-excitability is triggered through a unique signalling mechanism.

Axon injury induces calcium efflux from ER through IP3Rs and RyRs as well as calcium influx through the plasma membrane leading to elevation of intra-axonal calcium levels. Thus, perturbation of calcium homeostasis and ER calcium depletion following axon injury stimulates axonal ER stress. Importantly, inhibiting ER-mediated stress signaling in cerebral ischemia rat models has neuroprotective effects (Li et al. 2005). Inhibiting RyRs with Dantrolene reduced neuronal injury in an ischemic gerbil model (Wei and Perry 1996). Injury also causes decreased ATP production, which leads to inefficient protein folding (misfolded proteins) and ER stress (Roussel et al. 2013)(Li et al. 2013). ER stress may cause the unfolded protein response (UPR), a signaling program that can either promote cell survival or initiate a series of events leading to cell death. Canonical UPR stress sensor proteins, protein kinase RNA-like ER kinase (PERK), inositol-requiring protein-1 (IRE1α), and activating transcription factor-6 (ATF6), all bind to the ER chaperone immunoglobulin-binding protein (BiP). When ER calcium levels are depleted PERK activates calcineurin, which in turn dephosphorylates calnexin restoring ER calcium by activating SERCA (Wang et al. 2013). Increased SERCA expression in the injured neuron (Fig. 5) might help maintain calcium homeostasis.

The downstream signalling required for trans-synaptic communication remains to be elucidated. In peripheral neurons, locally initiated calcium waves can propagate to the nucleus to induce a transcriptional response (Cho et al. 2013). Other studies suggest that local influx of calcium may be a priming effect for retrograde microtubule-based transport of signaling complexes required to initiate transcription (Rishal and Fainzilber 2014). The increase of unfolded/misfolded proteins recruits more BiP, preventing its binding to sensor proteins. Without BiP, PERK dimerizes and autophosphylates. PERK phosphorylates eukaryotic translation initiation factor 2α (eIF2α) which causes inhibition of cap-dependent mRNA translation. Inhibiting PERK signaling via the small molecule ISRIB improved learning and reversed memory deficits following traumatic brain injury (Sidrauski et al. 2013; Chou et al. 2017). Phosphorylation of eIF2α increases translation through the internal ribosomal entry site (IRES) of ATF4. Interestingly, ATF4 mRNA localizes to axons and has been shown to locally translate in response to pathological Aβ_1-42_ microenvironments at axons (Baleriola et al. 2014). Locally synthesized ATF4 may be retrogradely transported to the soma to cause somatic ER stress and associated differential gene expression. Thus, an alternative to an axon injury-induced cytosolic calcium wave is a local calcium primed message involving retrograde transport.

We previously identified netrin-1, a known synaptogenic cue, as downregulated within the somatodendritic compartment both *in vivo* and *in vitro* following axon damage (Nagendran et al. 2017). Further application of exogenous netrin-1 after axotomy normalized both dendritic spine density and inhibitory terminals, suggesting netrin-1 signaling may regulate axotomy-induced synapse loss. Interestingly, ER stress activates IRE1, a sensor that activates UPR, to degrade netrin-1 mRNA (Binet et al. 2013).

ER stress is not only implicated in acute neural injuries, but also in multiple neurological disorders (Ozcan and Tabas 2012), including Alzheimer’s disease, Parkinson’s disease, multiple sclerosis, amyotrophic lateral sclerosis, and prion diseases. In these diseases, axon damage is a first site of pathology, suggesting signaling from the axon may be a key early event and clear target for future therapeutics.

## Conflict of Interest

A.M.T. is an inventor of the multi-compartment microfluidic device (US 7419822 B2, EPO 1581612, EPO 2719756) and is Chief Scientist and a Member of Xona Microfluidics, LLC. T.N. declares no competing interest.

## Author Contributions

T.N. designed experiments, acquired data, analysed date, and wrote the manuscript. A.M.T. designed experiments and wrote the manuscript.

## Funding

The authors received financial support from the National Institute of Mental Health (R42 MH097377), the National Institute of Neurological Disorders and Stroke (R41 NS108895), the American Heart Association (17GRNT33700108) and Xona Microfluidics, LLC. The content is solely the responsibility of the authors and does not necessarily represent the official views of the National Institutes of Health.

## Acknowledgments

Viral resources were supported by the GT3 Core Facility of the Salk Institute with funding from NIH-NCI CCSG: P30 014195, an NINDS R24 Core Grant and funding from NEI. Imaging was partially performed at the Neuroscience Center Microscopy Core Facility, supported, in part, by funding from the NIH-NINDS Neuroscience Center Support Grant P30 NS045892 and the NIH-NICHD Intellectual and Developmental Disabilities Research Center Support Grant U54 D079124.

## References

Baleriola, J., C. A. Walker, Y. Y. Jean, J. F. Crary, C. M. Troy, P. L. Nagy and U. Hengst (2014). “Axonally synthesized ATF4 transmits a neurodegenerative signal across brain regions.” Cell 158(5): 1159–1172.

Binet, F., G. Mawambo, N. Sitaras, N. Tetreault, E. Lapalme, S. Favret, A. Cerani, D. Leboeuf, S. Tremblay, F. Rezende, A. M. Juan, A. Stahl, J. S. Joyal, E. Milot, R. J. Kaufman, M. Guimond, T. E. Kennedy and P. Sapieha (2013). “Neuronal ER stress impedes myeloid-cell-induced vascular regeneration through IRE1alpha degradation of netrin-1.” Cell metabolism 17(3): 353–371.

Cho, Y., R. Sloutsky, K. M. Naegle and V. Cavalli (2013). “Injury-induced HDAC5 nuclear export is essential for axon regeneration.” Cell 155(4): 894–908.

Chou, A., K. Krukowski, T. Jopson, P. J. Zhu, M. Costa-Mattioli, P. Walter and S. Rosi (2017). “Inhibition of the integrated stress response reverses cognitive deficits after traumatic brain injury.” Proc Natl Acad Sci U S A 114(31): E6420–E6426.

Frost, S. B., C. L. Dunham, S. Barbay, D. Krizsan-Agbas, M. K. Winter, D. J. Guggenmos and R. J. Nudo (2015). “Output Properties of the Cortical Hindlimb Motor Area in Spinal Cord-Injured Rats.” Journal of neurotrauma 32(21): 1666–1673.

Gao, X., P. Deng, Z. C. Xu and J. Chen (2011). “Moderate traumatic brain injury causes acute dendritic and synaptic degeneration in the hippocampal dentate gyrus.” PloS one 6(9): e24566.

Ghosh, A., S. Peduzzi, M. Snyder, R. Schneider, M. Starkey and M. E. Schwab (2012). “Heterogeneous spine loss in layer 5 cortical neurons after spinal cord injury.” Cerebral cortex 22(6): 1309–1317.

Li, F., T. Hayashi, G. Jin, K. Deguchi, S. Nagotani, I. Nagano, M. Shoji, P. H. Chan and K. Abe (2005). “The protective effect of dantrolene on ischemic neuronal cell death is associated with reduced expression of endoplasmic reticulum stress markers.” Brain Res 1048(1-2): 59–68.

Li, S., L. Yang, M. E. Selzer and Y. Hu (2013). “Neuronal endoplasmic reticulum stress in axon injury and neurodegeneration.” Ann Neurol 74(6): 768–777.

McIlwain, D. L. and V. B. Hoke (2005). “The role of the cytoskeleton in cell body enlargement, increased nuclear eccentricity and chromatolysis in axotomized spinal motor neurons.” BMC neuroscience 6: 19.

Meijering, E., M. Jacob, J. C. Sarria, P. Steiner, H. Hirling and M. Unser (2004). “Design and validation of a tool for neurite tracing and analysis in fluorescence microscopy images.” Cytometry A 58(2): 167–176.

Nagendran, T., R. S. Larsen, R. L. Bigler, S. B. Frost, B. D. Philpot, R. J. Nudo and A. M. Taylor (2017). “Distal axotomy enhances retrograde presynaptic excitability onto injured pyramidal neurons via trans-synaptic signaling.” Nature Communications 8(1): 625.

Ozcan, L. and I. Tabas (2012). “Role of Endoplasmic Reticulum Stress in Metabolic Disease and Other Disorders.” Annual Review of Medicine 63: 317–328.

Persson, A. K., I. Kim, P. Zhao, M. Estacion, J. A. Black and S. G. Waxman (2013). “Sodium channels contribute to degeneration of dorsal root ganglion neurites induced by mitochondrial dysfunction in an in vitro model of axonal injury.” The Journal of neuroscience: the official journal of the Society for Neuroscience 33(49): 19250–19261.

Rishal, I. and M. Fainzilber (2014). “Axon-soma communication in neuronal injury.” Nature reviews. Neuroscience 15(1): 32–42.

Roussel, B. D., A. J. Kruppa, E. Miranda, D. C. Crowther, D. A. Lomas and S. J. Marciniak (2013). “Endoplasmic reticulum dysfunction in neurological disease.” Lancet Neurol 12(1): 105–118.

Sidrauski, C., D. Acosta-Alvear, A. Khoutorsky, P. Vedantham, B. R. Hearn, H. Li, K. Gamache, C. M. Gallagher, K. K. Ang, C. Wilson, V. Okreglak, A. Ashkenazi, B. Hann, K. Nader, M. R. Arkin, A. R. Renslo, N. Sonenberg and P. Walter (2013). “Pharmacological brake-release of mRNA translation enhances cognitive memory.” Elife 2: e00498.

Sun, L., J. Shay, M. McLoed, K. Roodhouse, S. H. Chung, C. M. Clark, J. K. Pirri, M. J. Alkema and C. V. Gabel (2014). “Neuronal regeneration in C. elegans requires subcellular calcium release by ryanodine receptor channels and can be enhanced by optogenetic stimulation.” J Neurosci 34(48): 15947–15956.

Taylor, A. M., M. Blurton-Jones, S. W. Rhee, D. H. Cribbs, C. W. Cotman and N. L. Jeon (2005). “A microfluidic culture platform for CNS axonal injury, regeneration and transport.” Nature Methods 2(8): 599–605.

Taylor, A. M., J. Wu, H. C. Tai and E. M. Schuman (2013). “Axonal translation of beta-catenin regulates synaptic vesicle dynamics.” The Journal of Neuroscience 33(13): 5584–5589.

Wang, R., B. C. McGrath, R. F. Kopp, M. W. Roe, X. Tang, G. Chen and D. R. Cavener (2013). “Insulin secretion and Ca2+ dynamics in beta-cells are regulated by PERK (EIF2AK3) in concert with calcineurin.” J Biol Chem 288(47): 33824–33836.

Wei, H. and D. C. Perry (1996). “Dantrolene is cytoprotective in two models of neuronal cell death.” J Neurochem 67(6): 2390–2398.

Wickersham, I. R., S. Finke, K. K. Conzelmann and E. M. Callaway (2007). “Retrograde neuronal tracing with a deletion-mutant rabies virus.” Nature methods 4(1): 47–49.

